# Cerebrovascular responses to O2-CO2 exchange ratio under brief breath-hold challenge in patients with chronic mild traumatic brain injury

**DOI:** 10.1101/2021.04.22.441010

**Authors:** Suk Tak Chan, Cora Ordway, Ronald J. Calvanio, Ferdinando S. Buonanno, Bruce R. Rosen, Kenneth K. Kwong

## Abstract

Breath-by-breath O_2_-CO_2_ exchange ratio (bER) is a respiratory gas exchange (RGE) metric, which is the ratio of the changes in the partial pressure of O_2_ (ΔPO_2_) to CO_2_ (ΔPCO_2_) between end-inspiration and end-expiration, has been demonstrated to characterize the cerebrovascular responses to breath-hold challenge in healthy individuals. We aimed to explore if bER could characterize cerebrovascular responses in patients with chronic mild traumatic brain injury (mTBI) under breath-hold challenge. We also investigated the correlation between bER and the severity of post-concussion symptoms.

Blood-oxygenation-level-dependent (BOLD) images were acquired using functional magnetic resonance imaging (fMRI) on eleven patients with chronic mTBI and eleven controls without brain injury history when performing a breath-hold task. Time series of RGE metrics of ΔPO_2_, ΔPCO_2_, and bER were computed, and their cross-correlation with regional change in BOLD (ΔBOLD) was calculated. Symptom burden was assessed using the Rivermead Post Concussion Questionnaire (RPQ), and its correlation with RGE changes was also measured.

Compared with controls, a diffuse decrease in the correlation between regional ΔBOLD and bER was found in the brain of mTBI patients (p_fdr_<0.05). No significant difference was found between patients and controls for the correlation of regional ΔBOLD with ΔPO_2_ and ΔPCO_2_. The averaged changes in bER (ρ=0.79, p=0.01) and ΔPO_2_ (ρ=0.70, p=0.03) in breath-hold epochs decreased with increased symptom severity indicated by RPQ scores.

Our imaging and symptom severity findings suggest that bER can be used to characterize cerebrovascular responses to breath-hold in mTBI patients. RGE may contribute to the post-concussive symptom severity.

## Introduction

Traumatic brain injury (TBI) is a global public health problem.^1^ A spectrum of post-concussion symptoms varying from headaches, fatigue, sleep, and cognitive disturbances may persist for months or years after TBI.^2^ Perturbations in microvascular flow are believed to precede neuronal dysfunction after TBI.^3–5^

Exogenous carbon dioxide (CO_2_) challenge in conjunction with functional magnetic resonance imaging (fMRI) measurement of blood-oxygenation-level-dependent (BOLD) signal in the brain has been used to show reduced/delayed cerebrovascular reactivity (CVR) in patients with chronic TBI.^6–8^ However, CO_2_ administration involves a complex gas delivery setup, and questions about potentially excessive vasodilation remain a concern for TBI patients at their acute and sub-acute stages when intracranial hemorrhage may be a risk factor. An alternative method for CVR assessment is a breathhold challenge.^9–14^ Increase in cerebral blood flow (CBF) is well established during breath-holding. While authors in the recent neuroimaging literature believe that CBF increase during breath-holding is due to hypercapnia only, ^-^ our team attributed CBF increase in breath-holding to the synergistic effect of both mild hypoxemia and mild hypercapnia due to the continual oxygen (O_2_) uptake and CO_2_ release in the closed breathing circuit during breath-holding.^18^

An increase in O_2_ uptake had been repeatedly demonstrated to be larger than an increase in CO_2_ release during breath-holding since 1946, resulting in a reduction in respiratory exchange ratio (RER), a ratio of CO_2_ to O_2_ at a steady state.^19–24^ The RER results in these studies are consistent with the findings in our recent breath-hold study, where the dynamic breath-by-breath O_2_-CO_2_ exchange ratio (bER), a ratio of breath-by-breath O_2_ uptake to CO_2_ release was increased during breath-holding.^18^ bER is defined as the ratio of the changes in the partial pressure of oxygen (ΔPO_2_) to the change in the partial pressure of carbon dioxide (ΔPCO_2_) between end-inspiration and end-expiration.^18^ The time-averaged value of bER is related to the reciprocal of RER. bER was found to demonstrate a strong positive correlation with dynamic global and regional cerebral hemodynamic responses under breath hold challenge;^18^ an increase in bER corresponds to an increase in CBF. bER changes were also superior to ΔPCO_2_ changes in coupling with global and regional cerebrovascular responses to breath-holding.^18^ The regional CVR in response to external CO_2_ challenge resembled that in response to bER, more than that in response to ΔPCO_2_ under breath hold challenge.^18^ Notably, ΔPCO_2_ is interchangeable with the end-tidal partial pressure of CO_2_ (P_ET_CO_2_) given the low ambient CO_2_ level. All these findings lend motivation to investigate if bER can characterize the cerebrovascular responses to breath-hold challenge in patients better than ΔPCO_2_ (P_ET_CO_2_).

In the current study, we aimed to characterize the cerebrovascular responses to RGE metrics of ΔPO_2_, ΔPCO_2_, and bER in patients with mild traumatic brain injury (mTBI) under breath-hold challenge. We hypothesized that the cerebrovascular responses to breath-hold challenge measured by BOLD signal changes (ΔBOLD) would have a stronger correlation with bER than with ΔPCO_2_ in patients with mTBI. ΔBOLD would also be less correlated with bER in compromised brain regions of patients than in normal brain regions of age-matched controls. Temporal and frequency features of RGE metrics in their coherence with ΔBOLD were also examined. The potential clinical impact was evaluated by correlating RGE metrics with symptom severity indicated by the Rivermead Post Concussion Questionnaire (RPQ) scores.

## Materials and Method

### Study design

We performed a prospective breath-hold fMRI study on a cohort of patients with chronic mTBI and age-matched controls without a history of brain injuries. All the experimental procedures were explained to the subjects. Signed informed consent was obtained before participating in the study. All procedures were performed in compliance with the Declaration of Helsinki and approved by the Massachusetts General Hospital (MGH) Human Research Committee.

### Participants

Patients diagnosed with mTBI for at least three months since the head impact were recruited at the MGH Neurology Clinic. The patients did not have a loss of consciousness or had a loss of consciousness of <30 minutes’ duration, and the Glasgow Coma Scale (GCS) score was >= 13.^25,26^ No abnormality was reported in the structural brain images. Control subjects without a history of brain injuries were also recruited.

### MRI Acquisition

Three-dimensional T1-MEMPRAGE (TR/TE1/TE2/TE3/TE4=2530/1.64/3.5/5.36/7.22ms, matrix=256×256×176, resolution=1×1×1mm^3^) and T2-SPACE FLAIR (TR/TE/TI=5000/355/1800ms, matrix=192×192×176, resolution=1×1×1mm^3^) images were acquired on a 3-Tesla scanner (Siemens Medical, Erlangen, Germany). BOLD-fMRI images were acquired with gradient-echo EPI sequence (TR/TE=1450/30ms, matrix=64×64, resolution=3.4×3.4×5mm^3^), while the subject was performing the breath-hold task. The timing of the breath-hold task was presented visually to the subject by a computer using the software Eprime Professional 2.0 (Psychology Software Tools, Inc., Pittsburgh, USA) (**Figure 1**). A breath-hold rehearsal was given before fMRI. No special instruction on the breath-holding after inhalation or exhalation was delivered to the subjects because BOLD signal amplitude has been suggested to depend on the depth of inspiration immediately preceding the breath-hold period,^27^ and end-expiration breath-hold may induce significant head motion.^28^

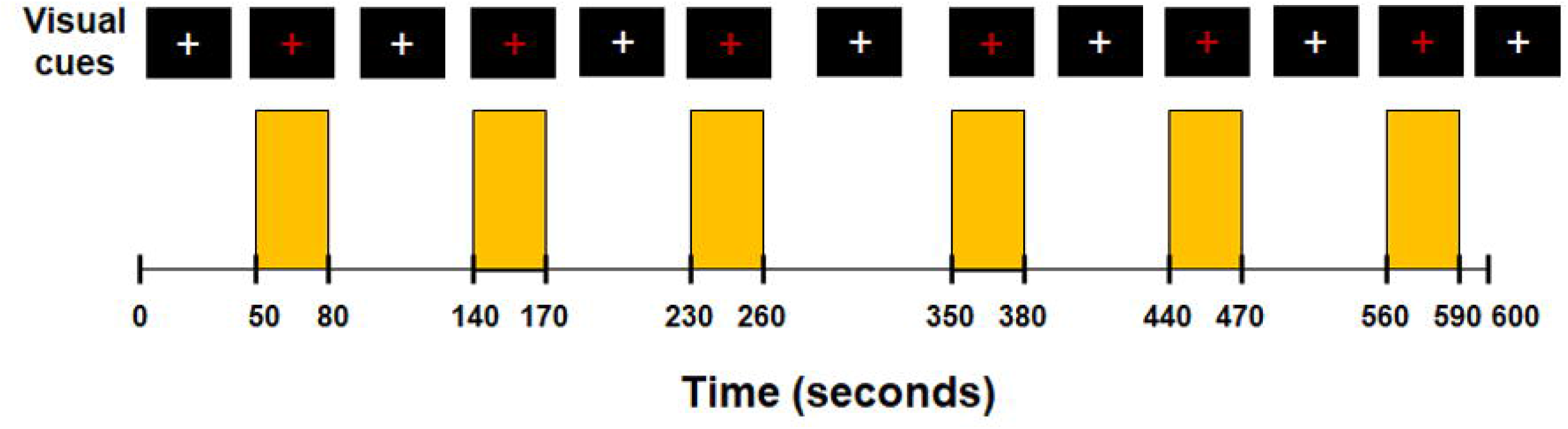

The partial pressure of O_2_ (PO_2_) and CO_2_ (PCO_2_) and respiration were measured simultaneously with MRI acquisition. A small nasal tubing was placed at the subject’s nostril to sample PCO_2_ and PO_2_ via gas analyzers (Capstar-100, Oxystar-100, CWE, Inc., PA, USA) after calibrating to the barometric pressure immediately before the scan and correcting for vapor pressure. Respiration was measured using a respiratory bellow connected to a spirometer (FE141, ADInstruments, Inc., CO, USA). All the physiological measurements were synchronized using trigger signals from the MRI scanner.

### Physiological data processing

The physiological data were analyzed using Matlab R2014a (Mathworks, Inc., Natick, MA, USA). Technical delay of PCO_2_ and PO_2_ was corrected by cross-correlating the time series of PCO_2_ and PO_2_ with respiratory phases determined from the respiratory bellow. End-inspiration (I) and end-expiration (E) were defined on the time series of PO_2_ and PCO_2_, and were verified by the inspiratory and expiratory phases on the respiration time series. Changes in the gas parameters in the breath-hold periods were interpolated by the values measured immediately before and after the breath-hold periods. Breath-by-breath O_2_-CO_2_ exchange ratio (bER) is defined as the ratio of the change in PO_2_ (ΔPO_2_=inspired PO_2_–expired PO_2_) to the change in PCO_2_ (ΔPCO_2_=expired PCO_2_–inspired PCO_2_) measured between end-inspiration and end-expiration in each respiratory cycle. The bER, ΔPO_2_ and ΔPCO_2_ measured in the baseline periods, i.e., the period before the first breath-hold epoch, were calculated. As the breath-hold duration might vary from one epoch to another, the changes of bER, ΔPO_2_ and ΔPCO_2_ in each breath-hold epoch were normalized to a 30-second breath-hold epoch by dividing the changes to the actual breath-hold duration in seconds and multiplying by 30.^21^ The normalized changes of each parameter over the six breath-hold epochs were then averaged.

### Image processing

All the BOLD-fMRI data were analyzed using software AFNI.^29^ The first 12 volumes of each functional dataset collected before equilibrium magnetization was reached, were discarded. Each functional dataset was corrected for slice timing, motion-corrected, and co-registered to the first image of the functional dataset. The time series of each voxel was detrended with the fifth order of polynomials to remove the low drift frequency, and the components of motion were removed using orthogonal projection. Head motions were quantified and compared between patients and controls (**Supplementary Figure 1**). The clean time series was then normalized to its mean intensity value. Voxels located within the ventricles and outside the brain were excluded from the following analyses of functional images. Individual subject brain volumes with time series of percent BOLD signal changes (ΔBOLD) were derived.

Hilbert Transform analysis^30^ was used to measure the cross-correlation between the ΔBOLD and the reference time series of RGE metrics (bER, ΔPO_2_ and ΔPCO_2_) in each voxel. Fisher’s Z-transformation was used to convert the cross-correlation coefficients in the individual subject maps to Fisher’s z scores. Individual subject brain volumes with Fisher’s z scores were then registered onto the subjects’ anatomical scans and transformed to the standardized space of Talairach and Tournoux^31^ for group analysis.

Wavelet Transform coherence^32^ was used to measure the dynamic changes in coupling between the time series of RGE metrics and ΔBOLD extracted from brain regions in the time-frequency domain. The analysis procedures were the same as those described in our other breath-hold study.^18^ The brain regions in gray and white matter were segmented using the software FreeSurfer.^33,34^ Example of squared wavelet coherence between bER and ΔBOLD in gray matter from a representative patient under breath-hold challenge is shown in **Supplementary Figure 2**. The percentage of significant coherence (sCoh%) defined as the percentage of time during which the magnitude of wavelet transform coherence was significantly different from the red noise outside the cone of influence was used to quantify the dynamic coupling between the time series of each RGE metric and ΔBOLD (**Supplementary Figure 2C**).

### Statistical analysis

Continuous variables (i.e., age and RGE metrics) were summarized as median and interquartile range, and group (mTBI and control) differences were assessed using the Kruskal-Wallis tests. Categorical variable (i.e., gender) was summarized as frequencies; group difference was assessed using the Chi-square test. Spearman correlation analyses were used to correlate in pairs the RPQ scores, age, and breath-hold duration and normalized changes in RGE metrics (bER, ΔPO_2_ and ΔPCO_2_) averaged over the six breath-hold epochs. Significant differences or correlations were considered at p<0.05.

Unpaired t-tests (3dttest++) in AFNI were used to compare the cerebrovascular responses to breath-hold challenge (i.e., Fisher’s z scores from Hilbert Transform analysis) between patients and controls. Multiple comparisons were corrected using equitable thresholding and clustering (ETAC) method.^35^ The group results were painted onto the averaged brain surfaces reconstructed by FreeSurfer.^33,34^

The sCoh% at the phase lags of 0±π/2 and π±π/2 from the individual subject wavelet transform analysis were separately plotted for mTBI and control groups to explore the Fourier periods/frequency bandwidths that oscillations of ΔBOLD in gray and white matter were in synchrony with each physiological time series of bER, ΔPO_2_ and ΔPCO_2_ when the subjects were performing breath-hold tasks. The sCoh% of RGE metrics with ΔBOLD in the six frequency bandwidths (0.008-0.015Hz, 0.016-0.03Hz, 0.031-0.062Hz, 0.063-0.124Hz, 0.125-0.249Hz, and 0.25-0.5Hz) extracted in the brain regions showing significant differences in Fisher’s z scores between mTBI and control groups was calculated for each subject. The distribution of sCoh% in these brain regions of the mTBI and control groups was then compared using the Kolmogorov-Smirnov test. Significant differences in the distribution of the sCoh% were considered at p<0.05.

## Results

Twenty subjects aged from 20 to 57 years were included. Ten of them were patients diagnosed with mTBI (median=38 years, interquartile range IQR=25.75-46.75 years; 5M and 5F). The selected clinical characteristics of the patients are shown in **Table 1**. The remaining ten were control subjects without brain injury history (median=28 years, IQR=25.25-34.25 years; 6M and 4F). No significant differences in age (p=0.34) and sex (p=0.65) were found between patients with mTBI and control subjects.

**Table 1.**
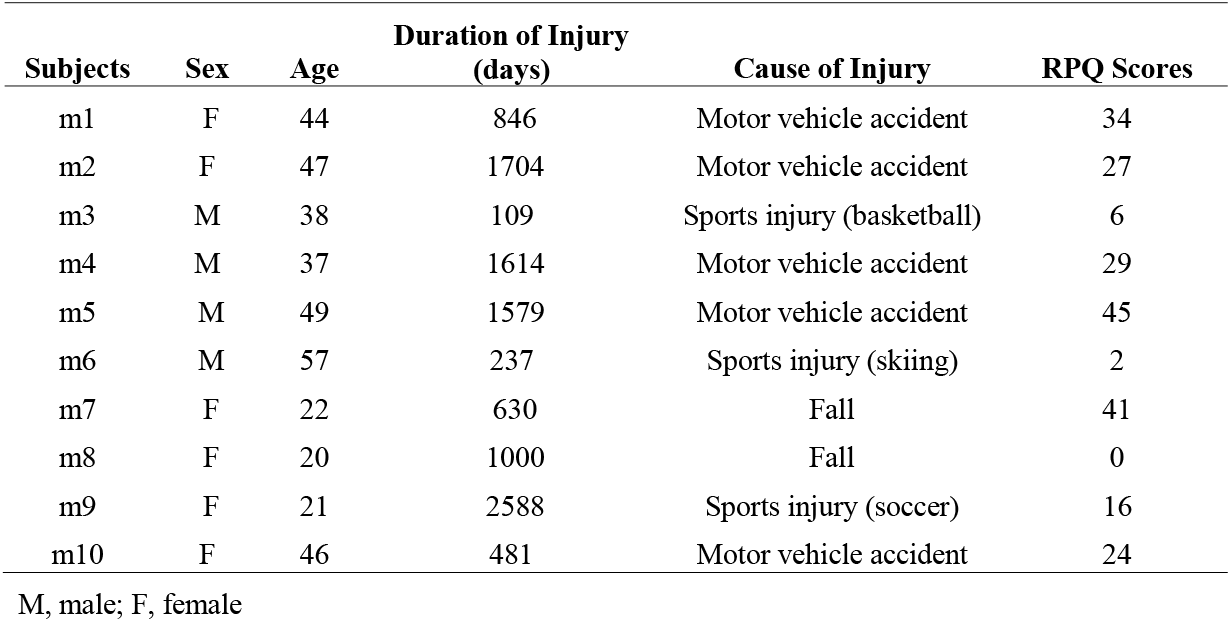
Selected clinical characteristics of the patients with mTBI.

The bER, ΔPO_2_ and ΔPCO_2_ measured in the baseline periods were summarized in **Table 2**. The averaged breath-hold duration, the averaged changes of bER, ΔPO_2_ and ΔPCO_2_ over the six breath-hold epochs were also included. The averaged changes of ΔPO_2_ during breath-holding were almost 3 to 4 folds larger than those of ΔPCO_2_. No significant difference in the RGE metrics was found between mTBI and control groups in the baseline periods and breath-hold epochs (p>0.05). No structural abnormality was identified in the T1-weighted and T2 FLAIR images of all subjects.

**Table 2.**
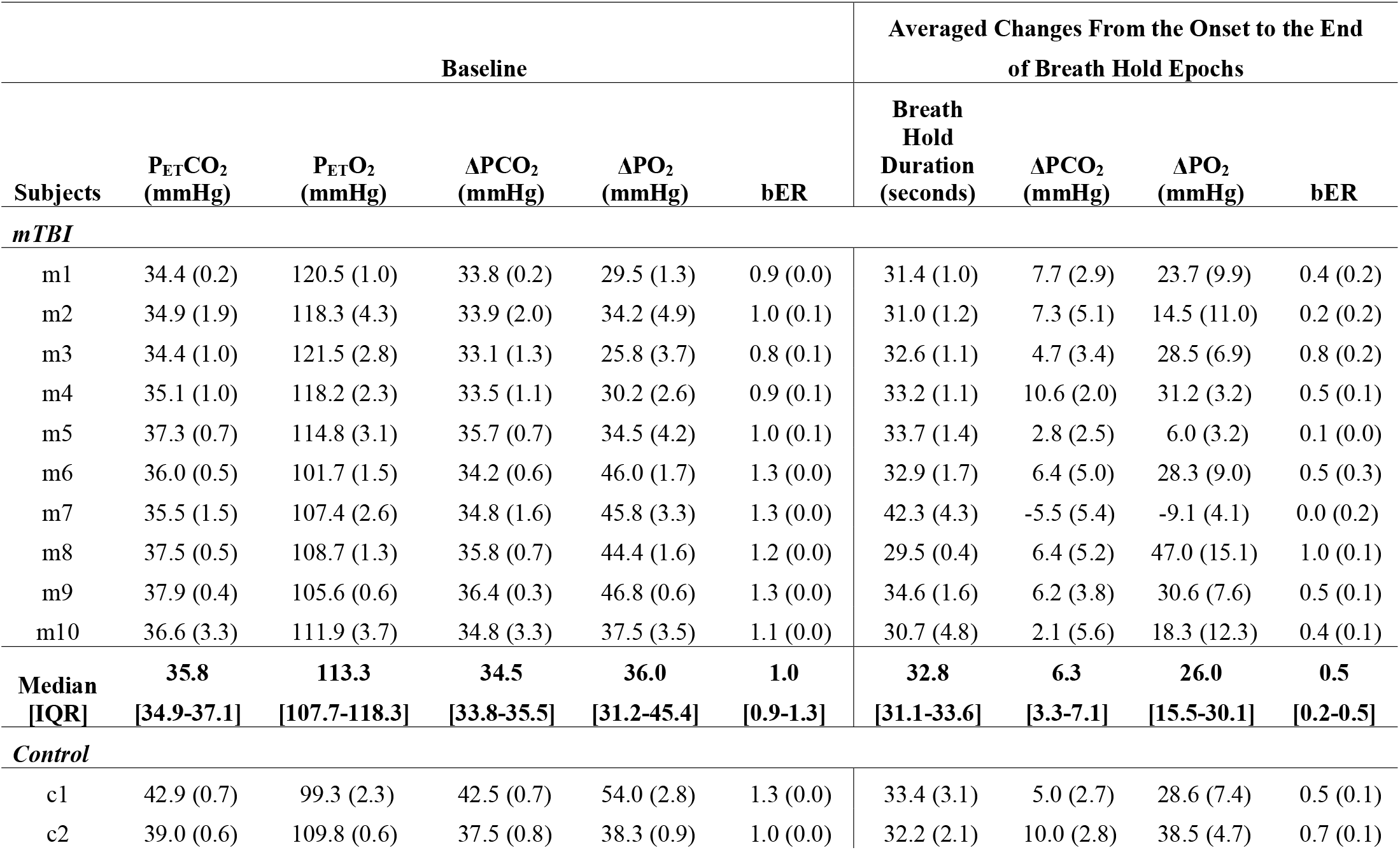
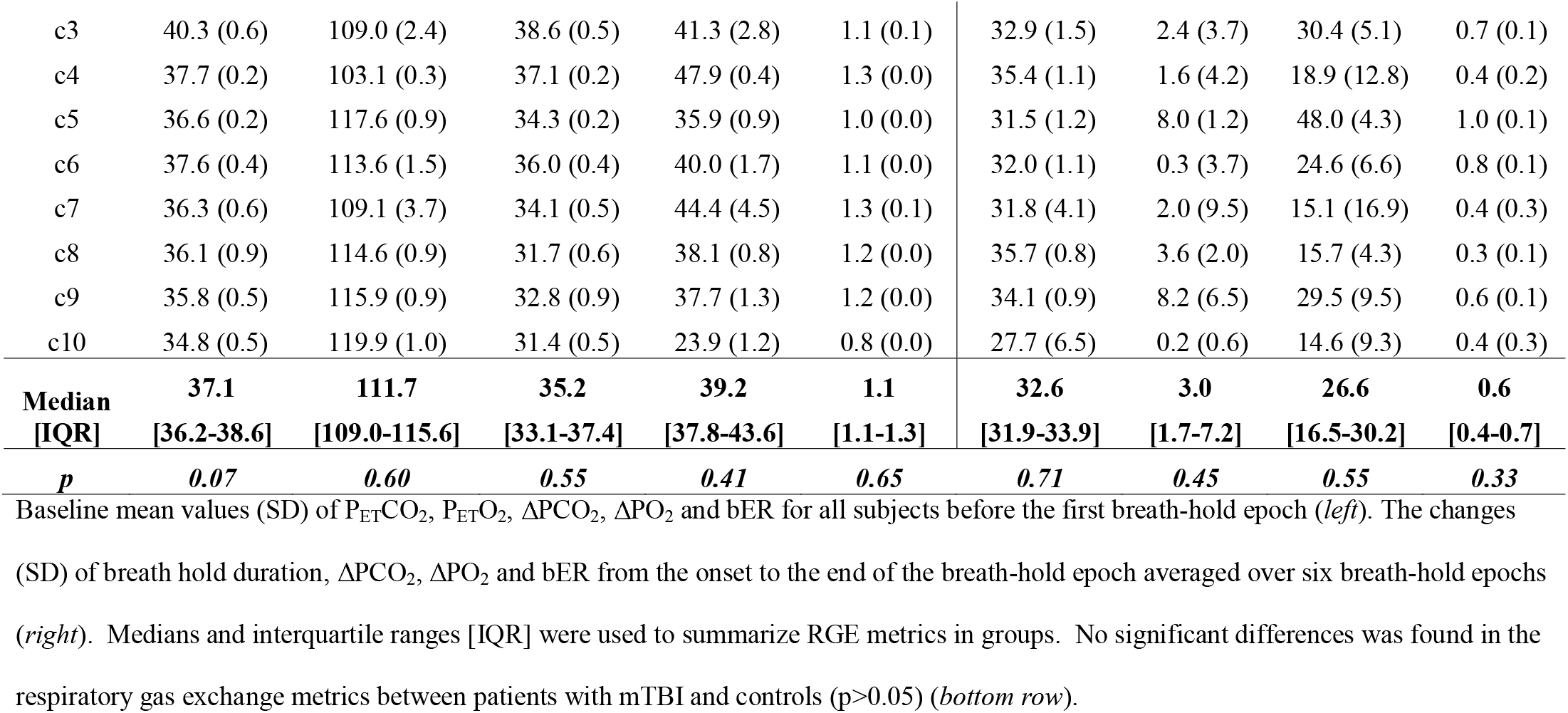
Baseline RGE values before the first breath-hold epoch and the averaged changes of RGE metrics during breath-holding.

### Correlation among RPQ scores, age and changes in RGE metrics

The correlation results among RPQ scores, age, breath-hold durations, and RGE metrics changes are summarized in **Figure 2**. During breath-holding, a significant correlation was found between RPQ scores and the averaged changes in bER (Spearman’s rho=-0.79, p=0.01), between RPQ scores and the averaged changes in ΔPO_2_ (Spearman’s rho=-0.70, p=0.03), and between the averaged changes in bER and ΔPO_2_ (Spearman’s rho=0.95, p<0.001). No significant correlation was found in the remaining correlation pairs (p>0.05).

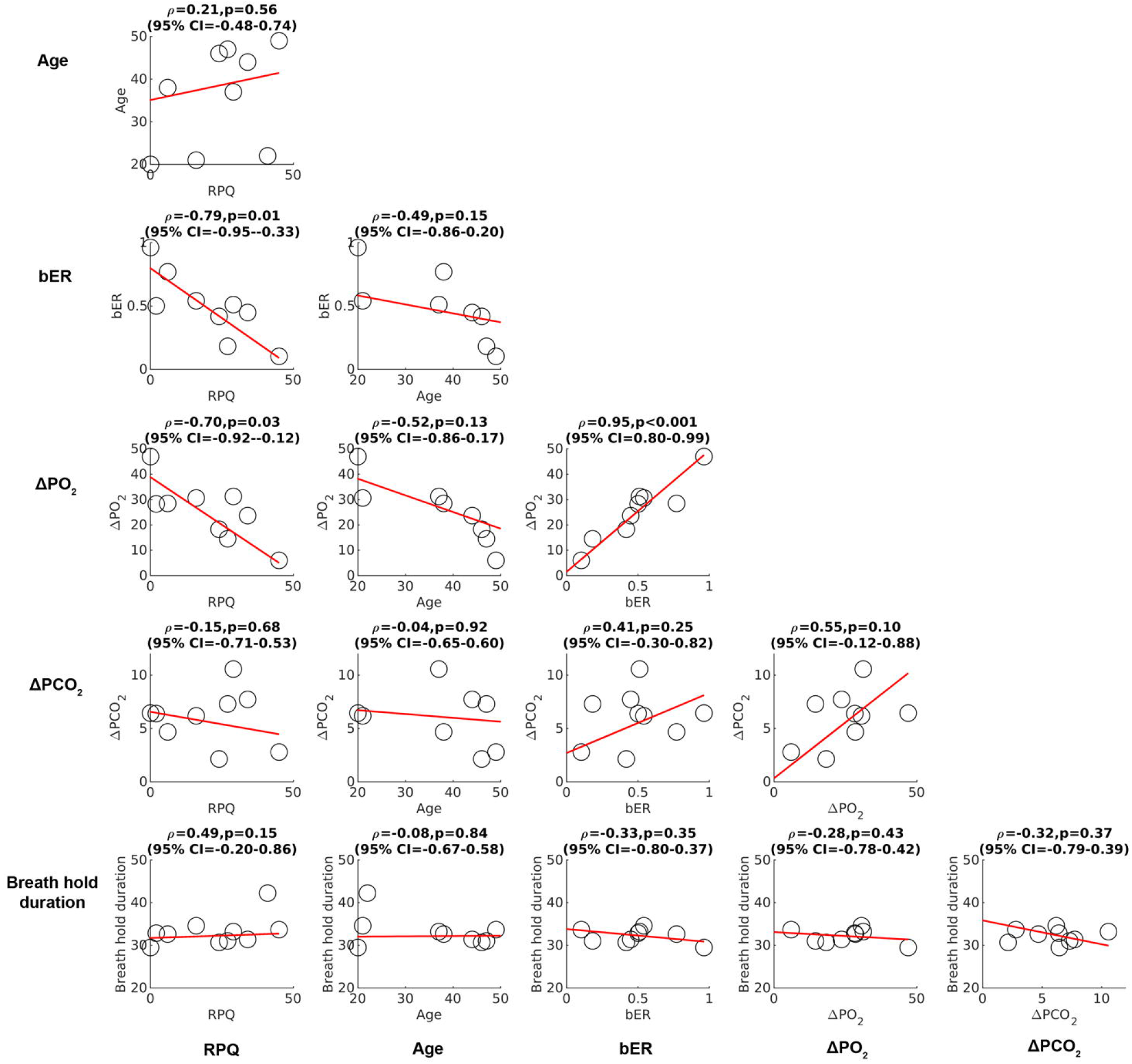

### Comparison of the cross-correlation of ΔBOLD with RGE metrics between patients with mTBI and controls

Time series of ΔBOLD in left gray matter and the corresponding RGE metrics in a representative mTBI patient and a control subject under breath-hold challenge are shown in **Figure 3**. The time series of bER followed more closely to ΔBOLD than ΔPO_2_ and ΔPCO_2_ in both representative patient and control subject.

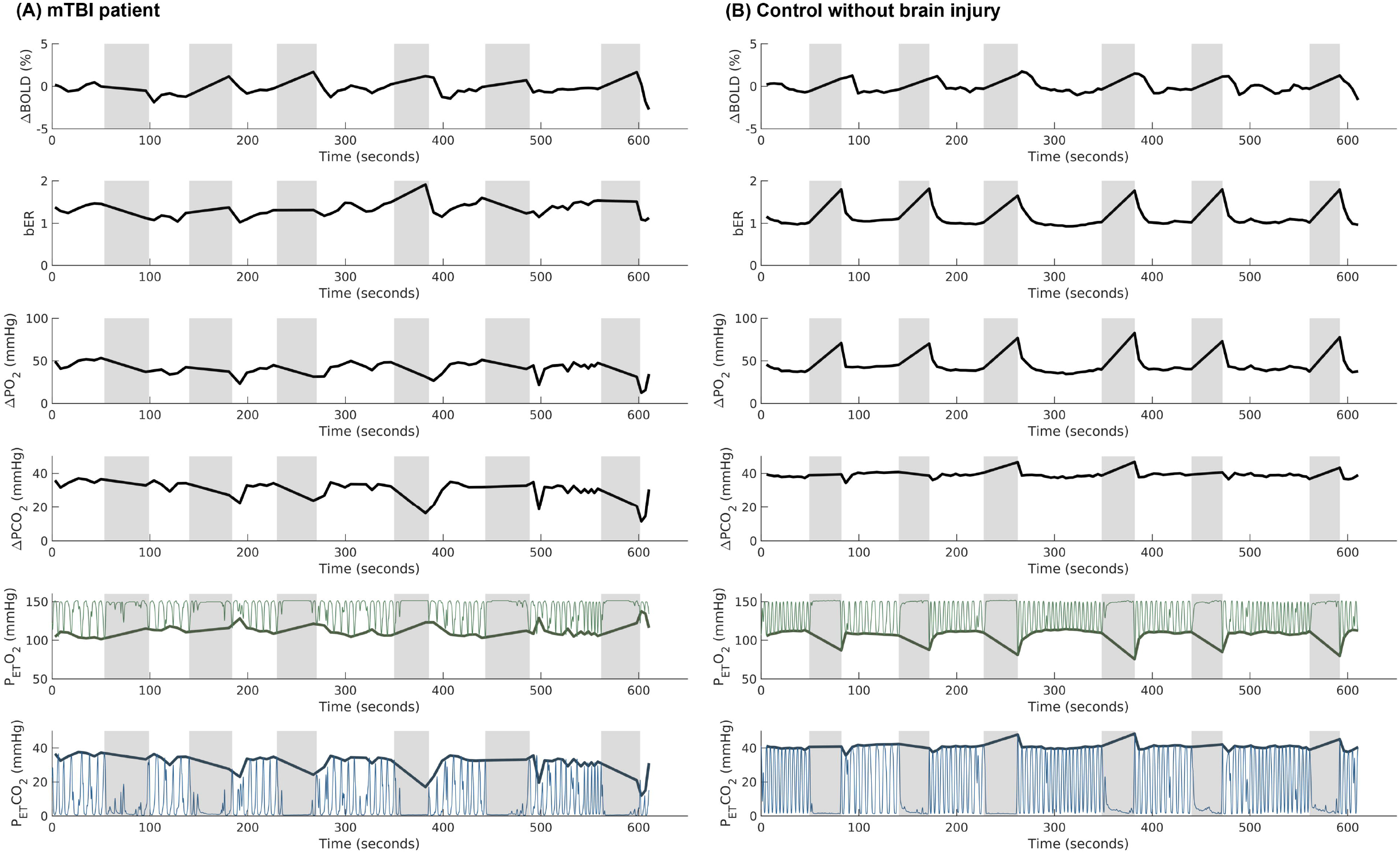

Compared with control subjects, a significant decrease in the cross-correlation between ΔBOLD and bER was mapped by fMRI in patients with mTBI at superior and middle frontal, precentral and postcentral, insular, superior parietal, middle and inferior temporal, paracentral, precuneus, posterior cingulate, cuneus, lingual, hippocampal and parahippocampal areas (p_fdr_<0.05) (**Figure 4**). No significant difference was found in the cross-correlation of ΔBOLD with ΔPO_2_ and ΔPCO_2_ between patients and controls.

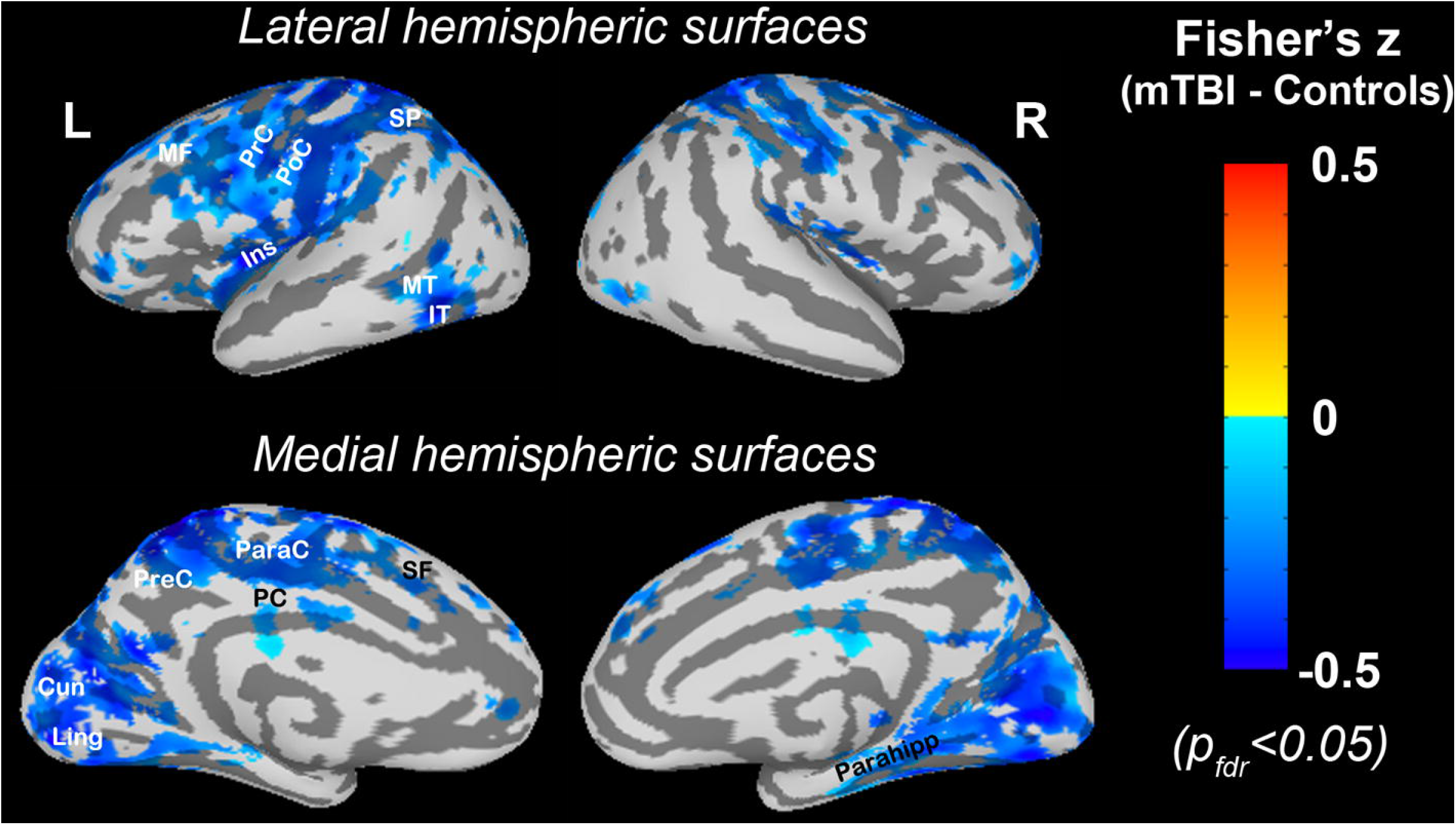

### Dynamic coupling between RGE metrics and ΔBOLD

In both the patients with mTBI and the control subjects, the dynamic coupling indicated by the sCoh% between bER and ΔBOLD at phase lag of 0±π/2 was the highest in the frequency range of 0.008-0.03 Hz while that between ΔPCO_2_ and ΔBOLD was the lowest in the same frequency range (**Figure 5**). The dynamic coupling between RGE metrics and ΔBOLD was weak at the phase lag of π±π/2 and the sCoh% stayed around 20% or below (**Supplementary Figure 3**). Comparing with control subjects, the dynamic coupling of bER and ΔPO_2_ with ΔBOLD in the frequency range of 0.008-0.03Hz and at phase lag of 0±π/2 was reduced in patients (**Figure 5**).

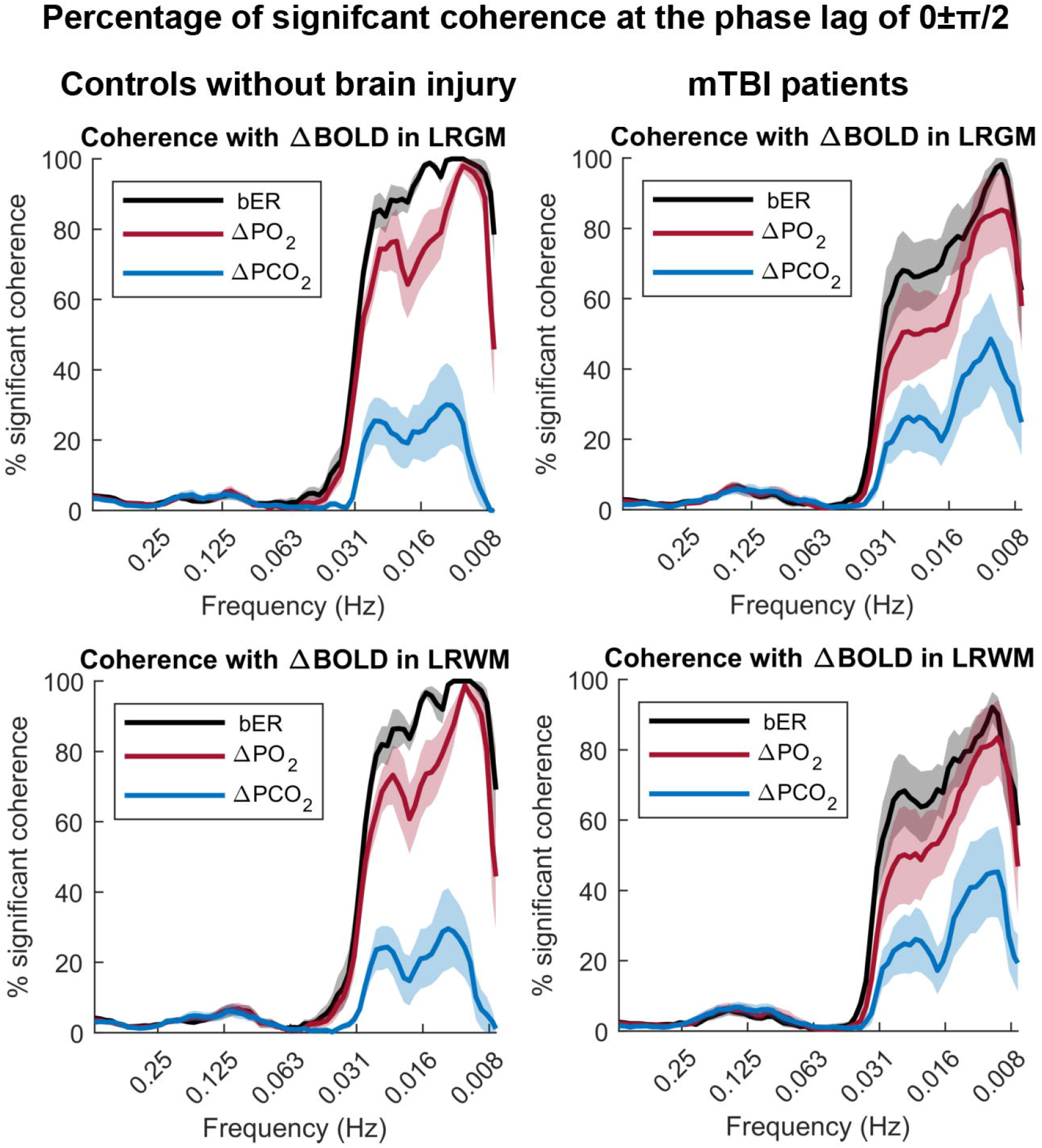

In the brain regions showing significant differences in the Fisher’s z scores between mTBI and control groups, the distribution of the sCoh% for the dynamic coupling of bER with ΔBOLD was significantly different between mTBI and control groups at the frequency bandwidths of 0.008-0.015Hz and phase lag of 0±π/2 (p=0.03) (**Figure 6**). No significant difference in the distribution of sCoh% was found between patients and controls for the dynamic coupling of ΔPO_2_ and ΔPCO_2_ with ΔBOLD in all frequency bandwidths and phase lag of 0±π/2.

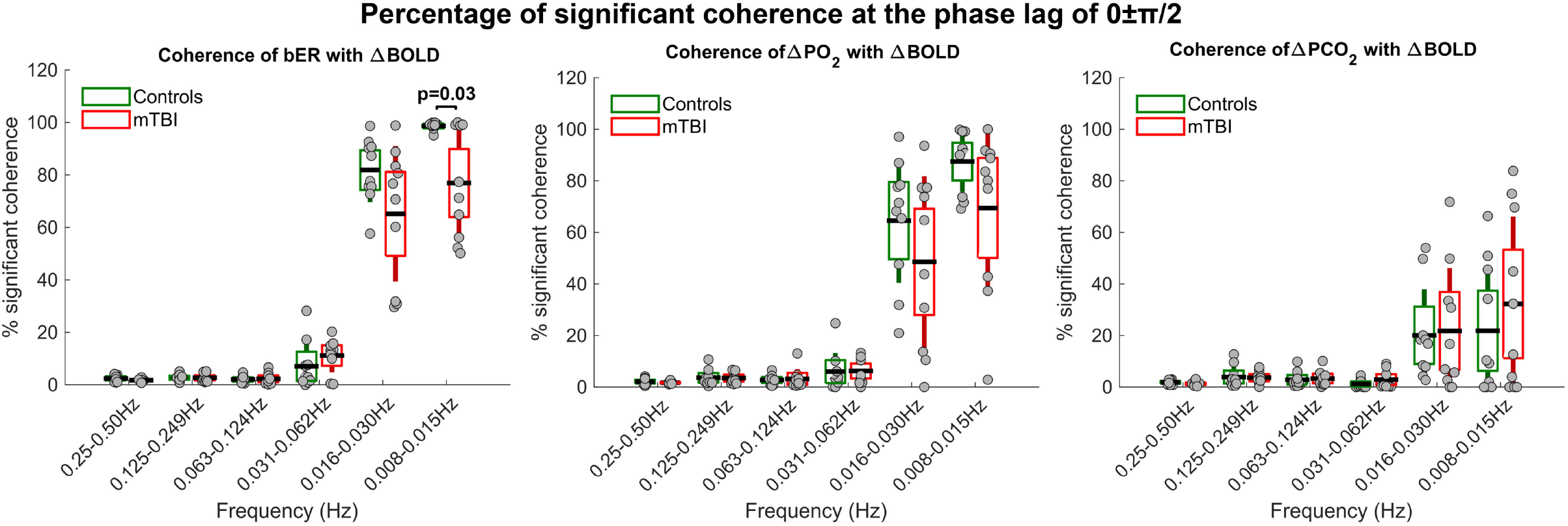

## Discussion

To our knowledge, this is the first study to characterize cerebrovascular responses with bER under brief breath-hold challenge in patients. We showed that the dynamic coupling of cerebrovascular responses with bER and ΔPO_2_ was stronger than that with ΔPCO_2_ in patients with chronic mTBI, similar to our previous breath hold results on healthy subjects.^18^ In patients with mTBI, the bER-ΔBOLD correlation under brief breath-hold challenge was significantly reduced in multiple brain regions when compared with control subjects who did not have a brain injury history. The averaged changes in bER and ΔPO_2_ in the breath-hold epochs decreased with increased symptom severity indicated by RPQ scores, while no such changes were observed between ΔPCO_2_ and RPQ scores. Patients with more severe symptoms showed a smaller increase in bER during breath-holding. The correlation between the symptom severity and the changes in bER and ΔPO_2_ during breath-holding could not be attributed to the breath-hold duration, as the changes in RGE metrics (bER, ΔPO_2_ and ΔPCO_2_) were normalized to a 30-second breath-hold epoch, and the breath-hold duration was not correlated with RPQ, age, or changes in RGE metrics.

Many research teams ^19–24^ including ours reported earlier a larger increase of O_2_ uptake than CO_2_ release under breath-hold challenge, resulting in reduced RER or increased bER for healthy subjects. Our current gas measurements on patients with mTBI again suggest the synergistic role of mild hypoxemia and mild hypercapnia during breath-holding. A strong correlation of the changes in bER with ΔPO_2_ but not with ΔPCO_2_ in patients during breath-holding (**Figure 2**) indicates that bER, being the ratio of ΔPO_2_ to ΔPCO_2_, is predominantly affected by ΔPO_2_.

In this study, we showed a significant decrease in the correlation between ΔBOLD and bER in response to breath-hold challenge in multiple brain regions of patients using Hilbert Transform analysis (**Figure 4**). These brain regions are responsible for the brain functions for attention, memory, visual processing, auditory processing, executive and autonomic functions.^36^ They also showed a general pattern that resembles the structurally atrophied regions of patients with mTBI reported by Warner et al.^36^ The intermittent cross-correlations between bER and ΔBOLD was then characterized as a function of both time and frequency in these brain regions using Wavelet Transform coherence analysis based on continuous wavelet transform.^32,37^

Like our previous breath-hold study in healthy individuals,^18^ bER had the most robust dynamic coupling with cerebral ΔBOLD for both mTBI patients and controls at the frequency range of 0.008-0.03Hz; ΔPCO_2_ had the weakest coupling while ΔPO_2_ had an intermediate coupling (**Figure 5**). Compared with controls, the coherence of bER with ΔBOLD was reduced in patients at the frequency range of 0.008 −0.03Hz. Concerns of using BOLD signal as a CBF surrogate had been addressed by another independent verification of the coupling between bER and CBF^18^ using CBF velocity measured by transcranial Doppler ultrasound at a sampling rate of 100Hz.

The dynamic coupling between bER and ΔBOLD was then measured in the selected brain regions showing a decreased correlation in Hilbert Transform analysis and showed significant intersubject variations among patients at the frequency range of 0.008-0.015Hz compared with controls (**Figure 6**). Such inter-subject variability may be due to the heterogeneity of brain injuries. bER, but not ΔPO_2_ or ΔPCO_2_, showed a significant difference in coupling with ΔBOLD between patients and controls at the frequency range of 0.008-0.015Hz. Given that bER is predominantly affected by ΔPO_2_, we attributed the somewhat surprising lack of statistical significance for ΔPO_2_ to the effect of ventilatory volume fluctuations common to both ΔPO_2_ and ΔPCO_2_. In comparison with ΔPO_2_, the format of bER as a ratio of ΔPO_2_ to ΔPCO_2_ has the advantage of being less affected by the ventilatory volume fluctuations resulting from isolated deep breaths. Future studies with a larger sample size are warranted to clarify the role between bER and ΔPO_2_.

## Conclusion

bER is a more useful metric to characterize the cerebrovascular responses to breath-hold challenge in patients with mTBI. The correlation between bER changes and symptom severity during breath-holding also suggests a relationship between RGE and post-concussion symptoms. Future studies with a large cohort of subjects would help explore the underlying mechanisms between RGE, cerebral hemodynamics, and post-concussion symptoms in mTBI patients.

## Authorship confirmation statement

| **Name** | **Location** | **Role** | **Contribution** |
| --- | --- | --- | --- |
| Suk Tak Chan, PhD | Massachusetts General<br>Hospital | Author | Design and conceptualized study; data collection and analysis; data interpretation; drafted the manuscript for intellectual content |
| Cora Ordway, BA | Massachusetts General<br>Hospital | Author | Data collection and analysis |
| Ronald J. Calvano, PhD | Massachusetts General<br>Hospital | Author | Design and conceptualized study; data analysis; data interpretation; revision of manuscript |
| Ferdinando S.<br>Buonanno, MD | Massachusetts General<br>Hospital | Author | Design and conceptualized study; data interpretation; revision of manuscript |
| Bruce R. Rosen, MD,<br>PhD | Massachusetts General<br>Hospital | Author | drafted the manuscript for intellectual content; revision of manuscript |
| Kenneth K. Kwong, PhD | Massachusetts General | Author | Design and |
|  | Hospital |  | conceptualized study;<br>data collection and<br>analysis; data<br>interpretation; drafted<br>the manuscript for<br>intellectual content;<br>revision of manuscript |

## Data availability statement

The original contributions presented in the study are included in the article/supplementary materials. Further inquiries can be directed to the corresponding author.

