## Supplementary figures and images for "Cerebrovascular responses to O2-CO2 exchange ratio under brief breath-hold challenge in patients with chronic mild traumatic brain injury"

### Supplementary Figure 1

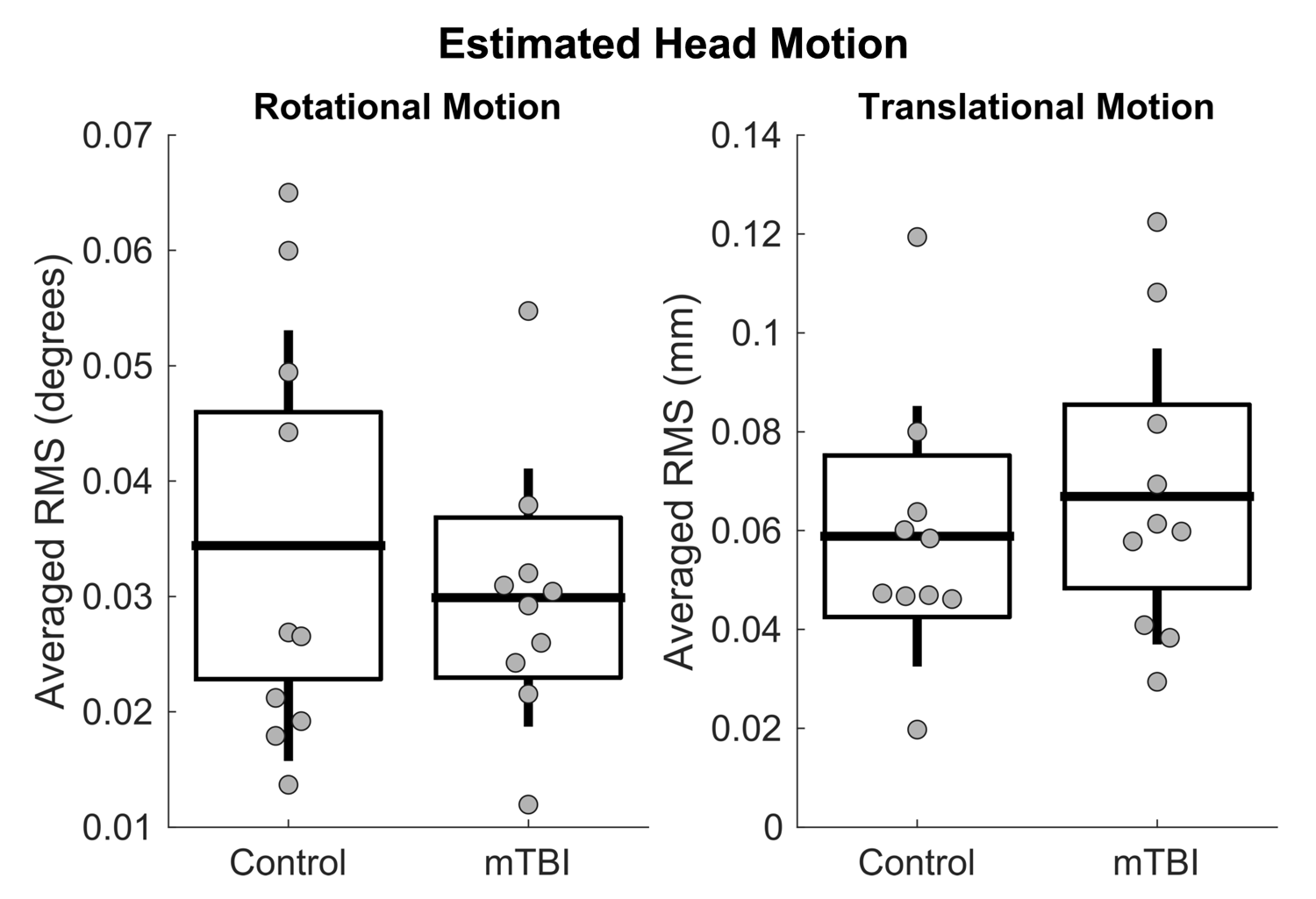

### Supplementary Figure 2

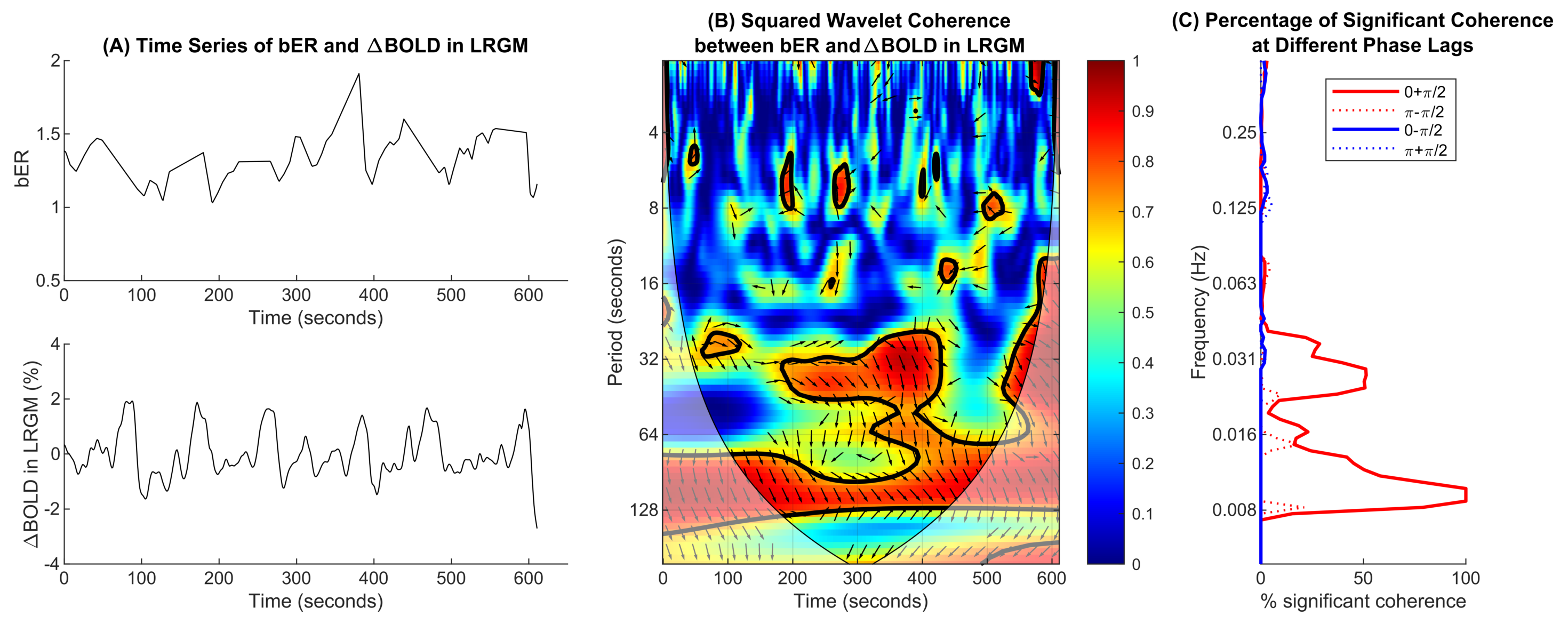

### Supplementary Figure 3

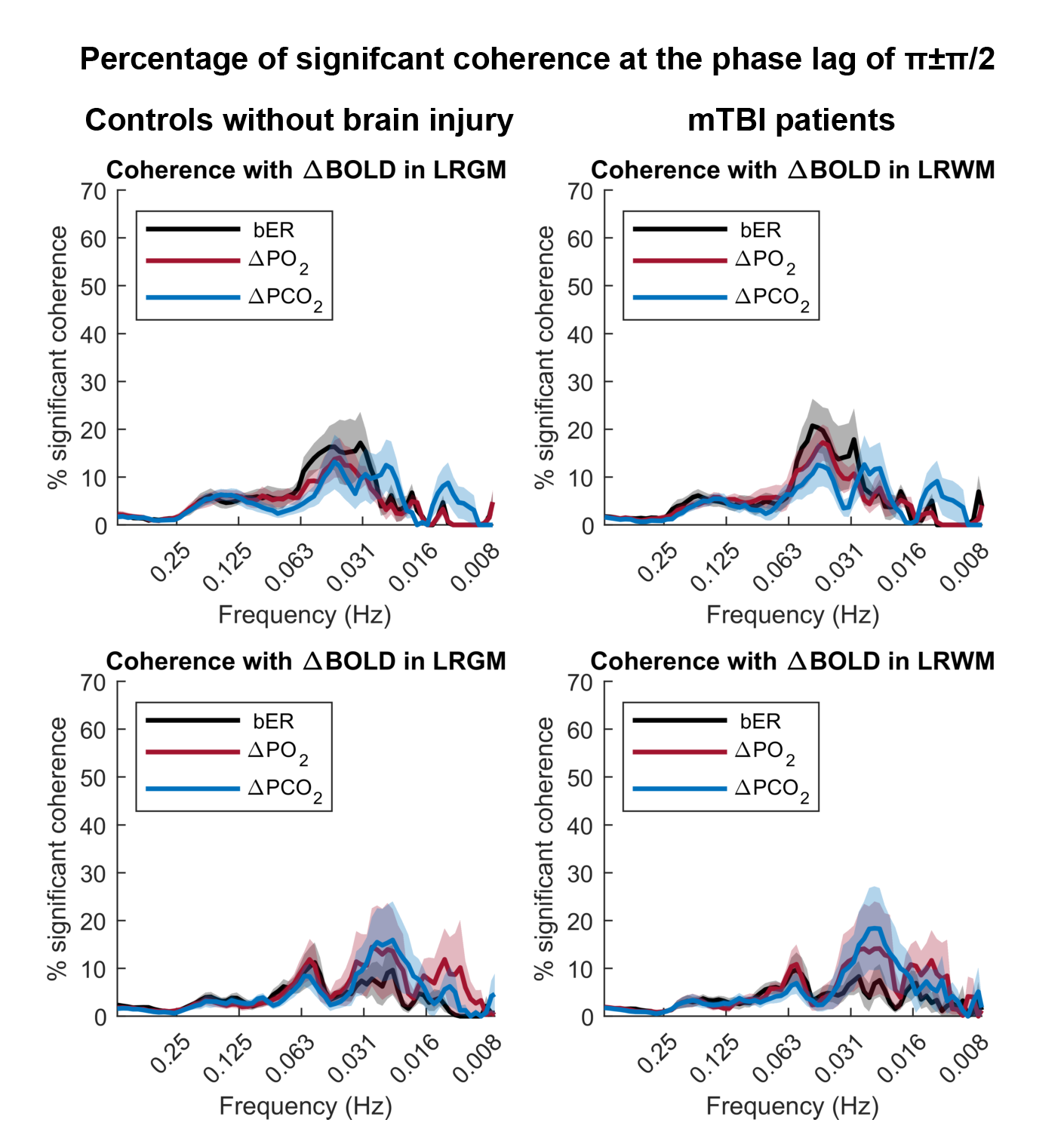
